# Knowledge, experiences and perceptions of the Ghana National Health Insurance Scheme in three districts

**DOI:** 10.1101/490359

**Authors:** Sataru Fuseini, Seddoh Anthony

## Abstract

**Background:** Ghana’s National Health Insurance Scheme is a demand side programme where the governing authority registers clients and purchases health care services for them from public and private providers. Access of services is high across a broad Benefits Package with no parallel enrolment necessary for any type of service at the point of access. Nonetheless, there is evidence of difficulty in acquiring and use of the NHIS card to access health care services.

**Objective:** While studies had been conducted into general awareness, there was no linkage between awareness, uptake and experiences with registration and use of the card. This study fills this gap.

**Methods:** This is a descriptive study. A mix of qualitative (39 Focus Group Discussions) and quantitative (625 household interviews) methods were used to collect the data. Qualitative data was analysed manually using a thematic approach while a frequency analysis was done for the quantitative data.

**Results:** Knowledge about the Scheme was near universal. Enrolment was lower among FGD discussants, 38.7% had valid cards, than for household respondents, 62.9% valid cards. While mixed experiences with the registration process was observed among FGD discussants, 74% of the households’ ranked attitudes of Scheme staff as positive. The study found the NHIS card facilitates access to facility based health care. Satisfaction levels with use of the card were mixed and contextual among discussants. However, 90% of households reported their cards were readily accepted at health facilities. Expired card (51.4%) and health facility had stopped accepting NHIS cards (14.3%) were mentioned as reasons for non-acceptance.

**Conclusion:** People’s experience during registration and use of the NHIS card to access health care has lasting effect on their perceptions of the Scheme. This can be harnessed to manage the high expectations, grow membership, discourage frivolous use and address artificial barriers of access.

## Introduction

From 1957, health care in Ghana was free until 1969 when the Hospital Fees Decree (NLCD 360) was passed, and subsequently amended as Hospital Fees Act 387 of 1970, to introduce user fee for consultation [1–4]. In the ensuing years, laboratory, and diagnostic services, invasive procedures and a select group of drugs were added to the chargeable list. Later, fixed fee charges for designated services and a 15% charge of the actual cost for general services was introduced with the enactment of The Hospitals Fees Legislative Instrument, LI 1313, in 1985 [3,4]. A list of exemptible services and conditions were specified in the legislation [1,3,4]. As the country sunk deeper into economic crisis, government funding for the health sector was reduced substantially, full cost recovery or what in Ghanaian parlance is referred to as ‘cash and carry’ came into effect in 1992 [4–8].

User fees persisted until 2005 when the National Health Insurance Scheme (NHIS) was fully rolled-out nationwide [9–13]. Building up to this roll-out was the passage in 2003 of Act 650 – the legal backing – which has since been replaced by Act 852 of 2012. The NHIS was a direct social response to the adverse effects of financial inaccessibility to health care services [5,11–13] for the over sixty-eight percent of the population at the time [14–15]. Act 650 established the National Health Insurance Authority (NHIA) – which has 10 regional offices and 159 District Mutual Health Insurance Schemes (DMHISs) – as the implementer of the NHIS. The DMHISs act as agency offices and are responsible for registration, card processing, revenue generation through premium collection, reimbursement of service providers and community engagement [9,16,17]. The Scheme is now widely recognised as a good pro-poor Social Health Insurance scheme [14,18].

About thirty-six percent (10,576,542 members) of the population is actively enrolled [19]. Designed as a demand-side programme, the NHIS run all year-round registration systems for clients and participating service providers [3]. Membership is compulsory for all persons living in Ghana based on an annual renewable system. This is however currently not being enforced [6,17,20,21]. At the point of registration, potential enrolees provide biometric information, which then serves as their identification within the system [16]. Enrolees are either premium-exempted or premium-paying. The premiums are set by the DMHISs, usually higher for urban and lower for rural populations, but within a range determined by the NHIA [6,17, 20,21]. An inclusion list of inpatient, outpatient and emergency services is covered at full cost by the curatively-inclined NHIS Benefits Package with no prior authorization before member access [3,22,23].

The NHIS has impacted positively on utilisation of outpatient (increased by more than forty-fold) and inpatient (increased by more than thirty-fold) services from the pre-NHIS era [24,25]. Whereas average per capita health care utilisation rate for NHIS cardholders is 3, that for general utilisation for Ghana is 1.7 [26,27]. Reports however shows of an ineffective registration system and unavailability of services to NHIS card holders [4,8,28]. Challenges of access to registration are related to both supply and demand side factors. While the supply side challenges are partly attributable to the switch from magnetic to biometric registration which has resulted in centralisation of the registration process at the district scheme offices, the demand side challenges are context specific [16–18,22,24]. Challenges of unavailability of services to cardholders can be categorised as NHIS systems-related failures, client abuse due to a lack of utilisation cap and general health system-wide failures. The NHIS systems-related failures though chronic, peaks and fluctuates. When normally otherwise NHIS clients are reported not to be receiving the full Benefits Package of services, at the peak of the Scheme’s challenges, service providers – mainly private and faith-based facilities – withhold all services to clients trying to access care with NHIS cards [4,8,18,29,30].

The effects of systemic challenges are easily describable from a technical angle. Their effects, however, from the populations view point are less clear and complicated. This necessitates the study of knowledge, experiences and perceptions that are context specific [31,32]. This study therefore aims to adduce and validate evidence that unpack issues that are reflective and necessary for gauging the perceived state of the Scheme by the population to improve its implementation. This study was undertaken as a baseline survey for the African Health Markets for Equity Programme (AHME). The NHIA had operated for over a decade. While studies had been conducted into general awareness, there was no linkage between awareness, uptake and experiences with registration and use of the card. This study set out to find; level of awareness; experiences with registration; usage after registration; and experiences with access to services.

## Methods

### Study Setting

The study was conducted in three districts: Ashaiman, Adaklu and Kassena-Nankana (divided as East and West). The districts were selected to represent the southern and northern sectors of the country and also the rural-urban divide.

Ashaiman, with an even sex distribution has a predominantly youthful population - 31.9% are between the ages 0-14 years - of 190,972 residents. The municipality is a bustling urban settlement in the Greater Accra region with 92% of the population economically active [33]. Accordingly, poverty prevalence is comparatively low at 4.4% [34]. 109,870 people were actively subscribed onto the Ashaiman Municipal Scheme [35].

Thirty-six percent of the 36,391 residents in the Adaklu district are below 15 years. Located in the Volta Region, this district exhibits features of rurality; 63.1% of the economically active population are engage in agriculture [36] resulting in 89.7% of the population living below the poverty line [34]. Sixty-seven percent active subscriptions were recorded by the district Scheme [35].

Split as East and West, Kassena-Nankana district is mainly rural (72.7%) and located in the Upper East Region. The population is 109,944 people. There are 19,790 households in the district averaging 5.4 persons per household [37]. Poverty prevalence is 37.3% [34]. A total of 93,965 people were actively subscribed onto the district scheme [35].

### Study Design

This is a descriptive study. Data was collected using a mix of qualitative and quantitative techniques. The study was in two parts; Focus Group Discussions (FGDs) using a semi structured open ended questionnaire to collect qualitative data and household survey using a structured tool to collect quantitative data. Earlier empirical studies exploring perceptions and experiences about the NHIS in Ghana used similar study design and methods [4–7,11,13,18,20].

### Sampling and Data Collection

Communities for the AHME project in each district were determined based on consultations with Staffs of the District Department of Social Welfare and DMHISs. Both the FGDs and household surveys were conducted in the same communities in each district except for Kassena-Nankana where only FGDs were conducted. These communities were purposively selected based on distance from the community - within the same community, less than 20 minutes travel time and more than 20 minutes travel time - to an NHIS registration centre and an NHIS accredited health facility. Five communities each in Ashaiman municipality, and Kassena-Nankan districts and ten in the Adaklu district were selected for the study. The sample, for both FGDs and household survey, was dependent on the availability of respondents. FGD discussants were selected based on the following criteria: women with Children Under 5; men 20 years and above who are decision makers in their homes; mixed group of opinion and traditional leaders; and a mixed group of adolescents. Criteria for inclusion into the household survey included: not previously a discussant in the FGDs; being the head or nominated by the head of household to respond; and age 18 years and above. For the household survey, a household sampling interval of ten was observed in Ashaiman municipality and that for Adaklu district was four.

A group of experienced consultants with expertise in health systems research, health insurance and public health were assembled to develop the study guides for both the FGDs and the household survey based on findings from a literature review. The FGDs were conducted by these same experts while data collection at the household level was done by researchers with experience in conducting national level surveys. The researchers were fluent in at least one of the dominant languages spoken in one of the study districts. The study in each district was preceded by a week of intensive communication and community durbars on the AHME programme. On the days of data collection, the town criers went round to mobilise the people.

Information was collected on respondents’ socio-demographic information, knowledge, experiences in acquiring and using the NHIS card to access health care services and attitudes of NHIS staff and health workers at registration centres and health facilities. Thirty-nine FGDs, thirteen in each district, involving 336 participants were conducted in all the three districts - 103 discussants in Ashaiman, 109 in Adaklu and 124 in Kassena-Nankana districts. On average, there were 12 discussants in each FGD session. Method for recording the FGDs was note taking and tape recording. Two household surveys were conducted in Ashaiman and Adaklu involving a total of 625 respondents, 309 and 316 respectively.

### Data Analysis

The qualitative data was analysed according to predetermined themes identified during the literature review. The actual analysis involved reading and manual coding of transcripts. The initial stages involved comparison of transcripts and field notes of FGD researchers to ensure validity of responses. After initial independent readings of the transcripts, the authors including two other researchers convened for the final analysis. The findings are presented as expressed quotes.

The quantitative data was analysed using the Frequency Analysis method. The data was first entered in Microsoft Excel and to control for data entry errors, there were three rounds of data entry verification and validation. Findings are presented as percentages in tables.

### Characteristics of Survey Respondents

There were more male FGDs discussants, 53.6%, than females. On the other hand, there were more female household survey respondents, 56.5%. More than half of both the FGD discussants and household survey respondents were between the age ranges 31-60 years and 26-60 years respectively. Whereas majority of FGD discussants, 43%, were engaged in agriculture and other agriculture-related businesses, majority of the household respondents were traders, 26.6%. About thirty percent each of FGD discussants and household respondents were educated to the Senior High School and Junior High School and levels.

## Results

### Membership

Seventy-two percent of the FGD discussants had ever enrolled onto the Scheme with 38.7% holding valid cards. The discussants who had never enrolled mentioned reasons to include lack of money to pay the premium, and registration fees and perceptions of poor attitude towards NHIS card holders by health care workers. Others narrated graphic examples of experiences that informed their decisions not to enrol as shown below:

> “……I went to hospital with my father and there were a lot of people. The nurse came and asked for those who did not have the NHIS card, called them out and treated them first. My father was treated later but he died. So I reasoned that if my father had not had the card, but had had money in his pocket, he would have been alive. So I decided not to have the card”.
>
> — [Adaklu FGD discussant].

> “I do not even think that there is any compelling reason for me to go get an NHIS Card. My wife went to the Hospital last month with her card and had to pay for her folder, spent a long time to get attended to, then was given a prescription that we had to struggle to buy the medicines…..it’s no longer attractive, and I think the challenges the NHIS faces in Bongo is too much”.
>
> — [Kassena-Nankana FGD discussant]

Two-fifth of the discussants who had the card had never renewed and some examples of the reasons for non-renewal are as below:

> “I was a valid card holder and went to the hospital but the machine could not read the card and I had to pay. When I compared access to health care with the card, I realized that not holding the card gives me better and faster access to health services”.
>
> — [Ashaiman FGD discussant]

> “When I fell sick and went to the hospital, I was asked to buy my own drugs, even though I had the card. So I bought all my medicines. I also hear other people with the same story. I felt it was not necessary to have the card. After all, health insurance does not cover all the drugs”.
>
> — [Adaklu FGD discussant]

> “I went to Bongo to get my NHIS card renewed, but the queue was too long; I spent three days and it was never my turn to be attended to…….they said the registration machine had broken down and there was no network, so I left in anger and never returned!”.
>
> — [Kassena-Nankana FGD discussant]

Seventy-eight percent of the household respondents had ever enrolled onto the Scheme and majority of them, 62.9%, had valid cards. As Table 1 shows, of the 22% that had never enrolled 56.1% stated they have not had the time to go register.

**Table 1:** Reasons for never enrolling onto the NHIS among household respondents.

| Reasons | Frequency (%)<br>(N=132) |
| --- | --- |
| Have not had time to register | 56.1 |
| Have not heard about the registration | 12.1 |
| Can pay for health care myself | 11.4 |
| Can't afford – too expensive | 12.9 |
| Insurance does not cover services I need | 16.7 |
| Hospitals don't respect the card | 3.8 |
| Processing procedures for card holders too long | 7.6 |
| Poor attitude of health workers towards card holders | 3.8 |
| Cheap medicines given to card holders | 7.6 |

### Knowledge about the NHIS

Knowledge about the existence of the NHIS was universal among the FGD discussants. They commonly understood and stated that the NHIS was implemented to facilitate the provision of *‘free health care’*. Some noted that the card facilitated access to hospital care without paying for the treatment. Others expressed how fee for service used to be catastrophic health expenditure for the household. A discussant at Kassena-Nankana indicated: *“we used to sell our cows and other food products to pay for the cost of hospitalization of our children when they get sick, but now with the NHIS card, we do not have to do that. You are taken care of free-of-charge”*. This sentiment was echoed severally and in different forms by discussants in all the districts.

The discussants mentioned both formal and informal channels as the sources through which they got information about the NHIS. Most indicated they get their information about the NHIS through the television and radio and from local announcements at the lorry station. For some, their knowledge and sources of information was based on their experiences with the Scheme. Interestingly, these discussants stated that they heard about the NHIS at the point of sickness as shown in the quote below:

> “…Me, I got to hear about the NHIS when I got sick in 2008 and was to be hospitalized; the doctor in Bongo Hospital asked me to go for the NHIS card so that I would not have to pay from my own pocket any time I got sick. He (the doctor) educated me about the NHIS, (and) then asked me to see a person who was in-charge of NHIS in the hospital to help me get enrolled”.
>
> — [Kassena—Nankana FGD discussant]

Among the households surveyed, knowledge of the existence of the NHIS was 98.2%. Respondents were presented with result statements in relation to membership and the exemptions package and were required to assess their validity or otherwise as shown in Table 2. Ninety percent (90%) correctly stated that renewal of membership onto the Scheme is on annual basis. Further, 54.7% correctly stated that pregnant women are enrolled onto the Scheme for free. Respondents’ knowledge about the Scheme’s benefit package was equally high. Over 80% of all respondents stated services covered by the Scheme included consultations and medicines. Respondents also mentioned picking a consultation card (69.1%), out-patient care (54.2%), admissions (50.6%) and laboratory investigation (37%) as the other services covered by the Scheme.

**Table 2:** Household respondents’ knowledge about the NHIS.

| Item | Statement | True % | False % | Do not know % | Status |
| --- | --- | --- | --- | --- | --- |
| (N = 625) |  |  |  |  |  |
| 1 | Foreigners who want to join NHIS have to pay in dollars | 12.2 | 34.8 | 53 | False |
| 2 | All children under 18 years do not need to register | 10.9 | 81 | 8.1 | False |
| 3 | Children under 18 are covered under the NHIS | 83.3 | 9.9 | 6.2 | True |
| 4 | Only the father is required to pay for NHIS | 6.7 | 85.6 | 7.7 | False |
| 5 | Pregnant women also pay to join the NHIS | 24.8 | 54.7 | 20.5 | False |
| 6 | Children who are not in school are required to pay for NHIS | 26.2 | 57.6 | 16.2 | False |
| 7 | The elderly, 70 years and above, are exempted from contributions | 77.1 | 10.1 | 12.8 | True |
| 8 | Only women above 70 years are exempted from contributions | 27.4 | 55.8 | 16.8 | False |
| 9 | NHIS is for a political party | 4.2 | 82.2 | 13.6 | False |
| 10 | The very poor (indigents) are covered for free under the NHIS | 42.2 | 35.2 | 22.6 | True |
| 11 | Cocoa farmers do not pay for NHIS | 5.5 | 48.4 | 46.1 | False |
| 12 | Rich people do not pay for NHIS | 3.8 | 86.7 | 9.4 | False |
| 13 | You renew your NHIS card every year | 89.9 | 2.1 | 8 | True |
| 14 | NHIS card is valid for five years | 81.6 | 5.9 | 12.5 | True |

### Experiences with the NHIS Registration and Card Processing

Responses of the FGD discussants provide insight into reasons for delays in registration and card processing. Discussants expressed frustration in the registration process. Some recalled experiences three days prior to the discussion to include persistent network failure and poor organisation of services at the district scheme offices. Two quotes from Kassena-Nankana below epitomise the general experience as narrated by the discussants.

> “Now all of us have to go to Kassena-Nankana in order to register for (the) NHIS card or renew our cards; it take(s) up to 5 days sometimes before you get to be attended to in Kassena-Nankana”.
>
> “When I went to register, the queue was so long that I had to sleep over there for 3 days before I got my NHIS card. I slept on a bench the first day and on the bare floor the second day. Getting food to eat was very difficult. They (DMHIS) did not even provide us with water to drink”.

However, majority of the household respondents got their cards the same day (56.3%) of registering. Others mentioned 2-5 days (12%), 1-2 weeks (9.1%), 2-4 weeks (4.1%), 1-3 months (17.7%) and three months and above (0.8%). DMHISs have autonomy in setting the fees for enrolment. Most of the enrolled respondents (87.8%) reported paying a fee of which 67.1% knew the fee amount before they went to register. Forty-one percent (40.9%) of these fee-paying registrants reported paying between – the exchange rate at the time – US$4.93-US$11.50 and 37.1% reported paying between US$2.58-US$4.69. Other amounts reported to have been paid before registration include less than US$1.04 (6.4%), US$1.04-US$2.35 (12.1%), US$11.74-US$18.78 (3.1%) and above US$18.78 (0.4%). Respondents ranked their impressions about the fees they paid as affordable (54.1%), moderately affordable (35.4%), high (8%) and too high (2.5%). Majority, 73.1%, thought the fees they paid were legal; 2.1% reported to have paid illegal fees and 20.8% stated they paid both legal and illegal fees to register. Of the respondents who did not pay a fee at the time of registering, 28.8% stated they had benefited from one of the exemptions policies.

### NHIS Staff Attitudes

FGD discussants that reported positive NHIS staff attitudes at registration centres were in the minority and from the Adaklu district. Discussants in the Ashaiman and Kassena-Nankana districts were unhappy with attitudes of some Scheme staff. The following quotes represent these sentiments:

> “The workers, especially those on attachment do not talk politely to clients at all. It is a very poor attitude. At times we go in to settle quarrels when commotion erupts during the registration process”.
>
> — [Ashaiman FGD discussant]

> “The NHIS staffs were disrespectful to us when we went to get registered in Kassena-Nankana. They insulted us by calling us animals. When we complained about the way they treated us unfairly by helping their friends and rich people to skip the queue they became angry and further insulted us….”.
>
> — [Kassena-Nankana FGD discussant]

Seventy-four percent (74%) of households’ surveyed ranked attitudes of NHIS staff at the point of registration as either very good or good and a further 17.1% ranked staff attitude as excellent. Eighty-three percent of respondents stated NHIS staff were patient and guided them through the registration process. However, 9% ranked the NHIS staff attitude as either poor or very poor. Reasons for the ranking of NHIS staff attitudes are presented in Table 3.

**Table 3:** Household respondents’ reasons for ranking of NHIS staff attitudes.

| Reasons | Frequency (%)<br>(N=493) |
| --- | --- |
| Patience and provided guidelines | 83.1 |
| Impatience and provided no guideline | 19.6 |
| No education of what NHIS is | 15.2 |
| Did not speak my language | 8 |
| Unfriendly and rude to me | 8.6 |
| Friendly and nice to me | 50.3 |
| Biased and favoured people they like | 11.6 |
| Allowed people to jump queue | 34.3 |
| They respected the queue | 36.8 |
| They have people to interpret and understood my language | 32.6 |
| They considered the elderly | 10.9 |
| They considered the poor | 3.4 |
| They discriminate against the poor | 0.8 |

### Access of Services and Experiences with Use of the NHIS Card at Health Facilities

Findings from the FGDs show that though the NHIS card is used for varied purposes, its common use is to access health care. Reasons mentioned by card holding discussants for their inability to use their NHIS cards to access health care are similar to that of the household respondents as presented below. Discussants who were satisfied with use of the card stated they did not pay for medicines and beds while on admission. They were given equal attention as the fee paying clients by health workers with some stating preferential treatment was rather given to card holders. In contrast, some stated that presenting the card at some health facilities was the reason for the poor quality of services that they received. Others also stated that use of the card was associated with some level of humiliation as shown in the following sentiments:

> “When you show up with the card at the hospital, the nurses look at you and complain that who is that poor person coming to disturb them”.
>
> — [Ashaiman FGD discussant]

> “My experience is that with the NHIS card you will not have attention compared with when you go without the card. I was the first to go and expected that I should be treated first before those who came after me. But non-cardholders who came after me were treated first”.
>
> — [Adaklu FGD discussant]

Seventy-three percent of household respondents that had used their NHIS cards reported using it to access health care. The frequency of use of the card ranged from less than 2 weeks (6.1%), 2-3 weeks (10.7%), 1-3 months (26.8%), 4-6 months (20.5%) and, 6 months or more prior to the survey (36%). The other common use of the card as found in this survey is as an Identity Card at the bank, Police station and to access mobile money services. On the other hand, majority of respondents who did not use their cards stated they had not fallen sick (88%) whilst holding a valid card. A quarter stated they had lost their cards. Other responses are provided in Table 4.

**Table 4:** Reasons for which household respondents had not used their NHIS cards to access care.

| Reasons | Frequency (%)<br>(N=100) |
| --- | --- |
| Never fallen sick | 88 |
| Lost the old card | 25 |
| Prefer Alternative and Traditional medicine | 20 |
| Can pay for health care myself | 15 |
| Insurance does not cover services I need | 2 |
| Card holders are treated last at the hospital | 4 |
| Hospitals don't respect the card | 1 |
| Waiting time to see the doctor too long | 3 |
| Processing procedures for card holders in the facility too long | 3 |
| Poor attitude of the health workers towards card holders | 3 |
| Cheap medicines given to card holders | 5 |
| Preference given to those who pay out-of-pocket | 1 |
| Card holders are asked to co-pay for health services | 8 |
| Others | 14 |

Seventy-two percent of the household respondents who had accessed care with their NHIS cards reported using it at a government health facility. Forty-six percent had used their cards at a private health facility. A further 14.8% had used the card at a pharmacy and the remaining 10.7% used their cards at faith-based, laboratory, and traditional (herbal) facilities. Private clinics (27.6%), district (18.8%) and regional (17.5%) hospitals were mentioned as the main places of last use of the card. On the last use, 37.6% of the household respondents stated they used it to access outpatient services. Thirty-three percent had used it to access pharmacy services. Respondents also mentioned having last used the card for admission (22.4%), laboratory (6.2%) and other services (0.9%). Ninety percent of the card holders who had used their cards stated health facilities easily and readily accepted their cards. Respondents whose cards were not easily and readily accepted mentioned the following reasons: card had expired (51.4%); health facility not NHIS accredited (25.7%); health facility had stopped accepting NHIS cards (14.3%); and others reasons (8.6%).

Majority, 53.6%, of the respondents whose cards were easily and readily accepted stated not all the services they required were covered by the NHIS card. Medicines (81.7%), laboratory services (57.5%), consultation (17.7%) and registration (17.2%) were mainly mentioned as not covered by the card. As a result, 53% reported paying additional money. Amounts mentioned to have been paid as additional fees included – exchange rate at the time – US$1.17-US$4.69 (35.3%), US$4.93-11.74 935.3%), US$11.97-US$23.47 (23.3%), US$23.71-35.21 (4.7%) and more than US$35.21 (5.3%). For those household respondents who reportedly paid additional monies, not all of them knew why they were charged the extra fees. The 83.9% of respondents who knew why they were charged extra fees mentioned the additional fees were mainly for medicines (38.2%) and laboratory services (36.9%). Further, 30% of the household respondents who had sought care stated they were referred to other facilities. Majority, 46.7%, stated they were referred to buy medicine. Other services mentioned for which respondents were referred included laboratory (26.7%), scan (15.6%), X-ray (10.4%) and other services (0.7%). Most of the respondents, 97%, who reported to have been referred stated they did not pay for services when they brought the results back.

Experiences with the use of the NHIS card to access health care among household respondents were mainly positive. Close to three-quarters of the about 77% of respondents that were either satisfied or very satisfied with use of their cards stated the health workers were friendly towards them. Despite the positive responses, there was some amount of dissatisfaction among some respondents with the services associated with the NHIS card as shown in Table 5.

**Table 5:** Reasons for household respondents ranking of their experiences of the use of the NHIS care to access care.

| Reasons | Frequency (%)<br>(N=365) |
| --- | --- |
| **Positive experiences** |  |
| Assurance that government will pay | 65.7 |
| Confident that I don't need to pay out-of-pocket to use services | 75.9 |
| As an NHIS card holder I was given preferential treatment | 59.4 |
| Staff attitude towards me (an NHIS card holder) was very friendly | 73.7 |
| Quality medication was made available to me (NHIS card holder) | 41.7 |
| Laboratory services was offered to me (NHIS card holder) freely | 5.3 |
| Others | 1.9 |
| **Negative experiences** |  |
| Waiting time to see the doctor too long | 51.4 |
| Processing procedures for card holders too long | 21.4 |
| Long queues at the OPD | 35.7 |
| Poor attitude of health workers | 40 |
| Doctor not present in the consulting room | 30 |
| No medicine available in the pharmacy | 28.6 |
| Laboratory services not working | 10 |
| Preference given to those who pay out-of-pocket | 21.4 |
| Staff attitude towards NHIS card holders is unfriendly | 25.7 |
| NHIS card holders are issued prescriptions to buy their own drugs | 42.9 |
| Card holders are asked to co-pay for health services | 37.1 |
| Others | 8.6 |

### Health Worker Attitudes towards NHIS Card Holders

Positive health worker attitudes toward card holders at the FGD level were mainly in the Adaklu district. *“The nurses insult the patients and their family for bringing people to the clinic if they are holding the NHIS card…..”* was a sentiment echoed severally and in different forms by most aggrieved discussants in Ashaiman and Kassena-Nankana districts as they recounted their negative experiences with health workers.

On the other hand, most household respondents (90.3%) reported of good, very good and excellent attitudes of health workers. Ten percent reported poor and very poor attitudes. Reasons for the respondents ranking of NHIS staff attitudes are presented in Table 6. Overall, 90.8% of respondents expressed a willingness to renew their cards. Reasons for renewal and non-renewal are shown in Table 7.

**Table 6:** Reasons for ranking of health worker attitudes by household respondents at health facilities.

| Reasons | Frequency (%)<br>(N=347) |
| --- | --- |
| Patience and provided guidelines | 64.4 |
| Impatience and provided no guideline | 11.2 |
| No education of what NHIS is | 2.7 |
| Did not speak my language | 6.4 |
| Unfriendly and rude to me | 26.4 |
| Friendly and nice to me | 52.7 |
| Biased and favoured people they like | 9.4 |
| Allowed people to jump queue | 43 |
| They respected the queue | 38.5 |
| They have people to interpret and understood my language | 43 |
| They considered the elderly | 18.2 |
| They considered the poor | 8.8 |
| They discriminate against the poor | 2.7 |
| They discriminate against the disable | 3.3 |

**Table 7:** Reasons for which household respondents will either renew or not renew their NHIS cards.

| Reasons for renewal (N=435) | Frequency (%) |
| --- | --- |
| Assurance that government will pay for the service | 20 |
| Confident that I don't need to pay out of pocket to use services | 25.3 |
| As an NHIS card holder I was given preferential treatment | 13.7 |
| Staff attitude towards me (an NHIS card holder) was very friendly | 15.3 |
| Quality medication was made available to me (NHIS card holder) | 12.7 |
| Laboratory services were offered to me (NHIS card holder) freely | 6.2 |
| Other, specify | 5.4 |
| **Reasons for non-renewal (N=44)** |  |
| Waiting time to see the doctor was too long | 11.7 |
| Processing procedures for card holders too long | 14.9 |
| Long queues at the OPD | 8.5 |
| Poor attitude of the health workers | 6.4 |
| Doctor not present in the consulting room | 6.4 |
| No medicines available in the pharmacy | 4.3 |
| Laboratory services not working | 2.1 |
| Preference given to those who pay out of pocket | 11.7 |
| Staff attitude towards NHIS card holders is unfriendly | 7.4 |
| NHIS card holders are issued prescriptions to buy their own drugs | 8.5 |
| Card holders are asked to co-pay for health services | 14.9 |
| Others, specify | 3.2 |

### Improving the NHIS

FGD discussants shared some ideas on how the NHIS could be improved and the changes they would like to see at the district scheme offices. Most discussants in all three districts would like to see the annual renewal of membership scrapped. They advocated for more flexible terms of membership beyond a year. Discussants also stated that card holders who had not used their cards before expiry should be given a discount on renewal. Others advocated for further decentralization of the registration process. They suggested district Scheme offices should open more registration centres to decongest the main offices to speed up the registration process. Majority of discussants also suggested that the district Scheme offices purchase more biometric registration equipment and find lasting solutions to the frequent system down-time and network failures. Others suggested as below:

> “The strategy applied to SSNIT (social security contributions) contributors should be expanded to include petty traders. Devise a mechanism to collect small amount of money periodically from them and register them onto the NHIS at very low tariffs”.
>
> — [Adaklu FGD discussant]

> “There is a need to increase the list of medicines that the NHIS covers especially for pain relief of women in labour. For now, as midwives working at the Health Centres and CHPS (Compounds), the patients and their relatives have to buy Pethidine or Diclofenac suppositories for pregnant women in labour or who are due for procedures such as episiotomy, post evacuation of the uterus, et cetera”.
>
> — [Kassena-Nankana FGD discussant]

## Discussion

This study sought to ascertain the knowledge, experiences and perceptions of the NHIS from the general populations’ perspective. An interesting finding was that over 70% of the study respondents had ever enrolled onto the Scheme. Enrolment was higher in Adaklu and Kassena-Nankana districts. These districts have high incidence of rural poverty. We therefore infer that respondents in these districts perceived the NHIS as protection from catastrophic health expenditures implying that the Scheme is fulfilling its core mandate [4,10,20,32,34]. Valid subscription was however 38.7% and 62.9% among FGD and household respondents. There appears to be high leakage or non-uptake of the Scheme. Solving this challenge of non-uptake and low renewal of membership has operational implications at the district Schemes level to invest resources to understand the causes and take remedial actions. Good social marketing of the Scheme at the community level in the meanwhile may contribute to membership growth and retention [6,8,29].

We observed that respondents had high knowledge of the NHIS. All FGD discussants and 98.2% of households surveyed were aware of the Scheme, and fairly informed on membership and the benefits package. This seemingly high knowledge may be attributable to the vigorous marketing in the early years [20] and the high political commitment exhibited towards the Scheme by successive governments [2,20,23]. Notwithstanding, the evidence was also indicative of some level of misunderstanding and misrepresentation of the Scheme. These findings are consistent with that reported by Gobah and Liang in a rural district of Ghana and Nelson et al in a metropolitan area in Southern Nigeria [20,38]. Continues engagement and communication between DMHISs and district level stakeholders will serve as a channel to disseminate timely, correct and critical information about the Scheme to communities. This will also serve to nurture a relationship of discourse and dialogue between district schemes and communities where some of these challenges faced by the Scheme can be addressed [29,32].

FGDs, as it is generally known, allows for greater self-expression hence the sharp contrasts in the narratives of experiences in obtaining the NHIS card. The introduction of a biometric system centralized the NHIS registration process. This has resulted in challenges of inadequate staff numbers, machines, and supplies and exacerbated by persistent system downtime and network failure. These are administrative bottlenecks that impact client experiences. However, for particular Schemes the challenge is mostly related to negative staff attitudes which were mostly referred to as poor. These challenges consequently affect enrolment and renewals. This brings into perspective the question on operational guidelines and efficiency of the card processing system at the district scheme offices [6,20]. The negative experiences associated with the NHIS on account of staff attitudes confirm earlier findings by Gobah and Liang and Jehu-Appiah et al [6,20]. This will require extensive training in crowd management and customer relations to solve [4,20,34].

From a methodological perspective, experiences with the NHIS registration process and attitude among the survey sample were mixed. We observed less favourable experiences among the FGD participants compared to the household respondents. Reflective discussions tend to draw from people narratives and higher levels of embellishment of lived experiences. As a result, experiences may become biased and exaggerated. To address this inherent bias, we tried to focus the groups on the issues as personal experiences rather than emphasising hypothetical sympathies with other participant experiences. This does not mean bias is totally eliminated. Human emotions once shared can become a power opinion manipulator and more so when the shared experience has a negative impact. We observed this in the FGD groups leading to discussants sharing their negative experiences more readily than in household individual surveys. Though contrasting, our interactions were more towards understanding all aspects of the lived experiences rather than finding a dominant pattern. Thus, as a descriptive study, the subjective opinions are as relevant as the perspective they bring to bear on the success of the health insurance scheme.

Once the NHIS card is acquired, it facilitated access to facility based care. This is evident in the high proportion of respondents who had used their cards when sick. These findings are therefore suggestive that not only does possession of a valid card reduce the barrier of financial access, it also promotes good health care seeking behaviour [6,9,10,20]. This does not mean that service experience was always satisfying as some had difficulty appreciating the quality they received. Dalingjong and Laar, Jehu-Appiah et al and Alfers observed similar experiences [6,7,23]. Though satisfaction levels with the use of the card were generally high, the negative experiences were symptomatic of the inefficiencies and deficiencies of the health system of Ghana [3,4,20]. These striking reactions in differences in the use of the card, we reiterate, provides a depth of evidence of what is working and what is not. Alhassan et al suggested that such negative reactions to services under the Scheme are attributable to the high expectations, which sometimes are unrealistic, that translate into demand [31].

At the provider level, increased utilization of health services in the aftermath of the introduction of the NHIS brought to the fore systemic challenges and the unpreparedness of the health service to implement such a policy. This resulted in overload of work at health facilities as health workers were not adequately trained to interact and process NHIS claims [3,6,23,]. The use of the NHIS card, however, among both sets of respondents depends on the individual need and acceptability. The primary use among all is to access care. It is the secondary use that shows variations.

We note some limitations of our study. An inherent limitation is that the sample size and sampling process for the household survey was not scientifically predetermined. Since we did not also focus on triangulating responses, there is the possibility of respondent recall bias. We note that strong views by vocal discussants during the FGDs may have influenced or drowned the views of the less vocal discussants. We do not also exclude the possibility of study respondents providing socially desirable answers when interviewed. That said, the use of both qualitative and quantitative methods allowed us to capture in the true words and weigh the knowledge, experiences and perceptions of the population [39] about the NHIS.

## Conclusion

This study has demonstrated that awareness and knowledge about the NHIS is universal. Re-enrolment onto the Scheme is however faltering. The use of the NHIS card to access health care is high. People’s experience during registration and use of the card has lasting effect on their perceptions of the Scheme. Formation of these perceptions is also dependent on the design of the Scheme, staff attitude and health system related factors. The Scheme factors relate to personnel, machines and supplies for registration. The service factors are mainly attitudinal, and the general organisation and service availability. That said, there are some immediate things that can be done to help. At its basic, the NHIA can ensure they have strong presence in communities by developing and maintaining communication channels for information dissemination and feedback. Secondly, District Offices should improve on their mobile registration centres and develop strategies to ease the long delays in registration. Staff should be trained in crowd management and SMS systems on information notification can help in prompting expiry dates and times of registration.

Overall, the experiences from this study about the Ghana NHIS contribute to improving the Scheme. The positive knowledge, experiences and perceptions can be harnessed by district schemes to manage the high expectations, grow membership and to discourage frivolous use of the card at service points. The negative experiences and reactions, on the other hand, if not sufficiently addressed can undermine the gains achieved and create artificial barriers of access to services under the Scheme.

## Abbreviations

AHME: African Health Markets for Equity
DMHIS: District Mutual Health Insurance Scheme
FGDs: Focus Group Discussion
NHIA: National Health Insurance Authority
NHIS: National Health Insurance Scheme

## Acknowledgements

We thank Cheremeh-Marfo Emmanuel for his contribution during the data collection and analysis process.

## Availability of data and materials

Data used in this analysis has been deposited at the African Health Markets for Equity (AHME) project data repository at https://ahmesocialdevtools.com/app/. Access can be granted upon request.

## Authors’ Contribution

All authors (SA & SF) conceptualized the study, collected, and analysed the data and drafted the manuscript. All authors read and approved the manuscript.

## Ethical Approval

The Research Divisions of the Ministry of Gender Children and Social Protection and the NHIA waived ethical approval for the AHME baseline survey as it was part of a larger piloted social intervention programme on identifying and registering poor people onto the NHIS for free. Informed consent, verbal, was obtained from all study participants. All identifiable traces linking respondents to statements have been removed.

## List of Figures

**Figure 1:** NHIS membership status of survey respondents

## References

1. Asenso-Okyere WK, Anum A, Osei-Akoto I, Adukonu A. Cost recovery in Ghana: are there any changes in health care seeking behaviour? Health Policy and Plannning. 1998;13(2):181–118.

2. Agyepong IA, Adjei S. Public social policy and development: a case study of Ghana National Health Insurance Scheme. Health Policy and Planning. 2007;23:150–160.

3. Seddoh A, Adjei S, and Nazzar A. Ghana’s National Health Insurance Scheme: Views on Progress, Observations, and Commentary. Accra: Rockefeller Foundation. Available from: http://www.ch-ghana.org/documents/Publication/Report%20on%20observations%20and%20commentary%20on%20NHIS.pdf.

4. Derbile EK, van der Geest S. Repackaging exemptions under National Health Insurance in Ghana: how can access for the poor be improved? Health Policy and Planning. 2013;28–586–595.

5. Blanchet NJ, Fink G, Osei-Okoto I. The Effects of Ghana’s National Health Insurance Scheme on Health Care Utilization. Ghana Medical Journal. 2012;46(No.2):76–84.

6. Jehu-Appiah C, Aryeetey G, Agyepong I, Spaan E, Baltussen R. Household perceptions and their implications for enrolment in the National Health Insurance Scheme of Ghana. Health Policy and Planning. 2011;27(3):1–12.

7. Dalinjon PA, Laar AS. The national health insurance scheme: perceptions and experiences of health care providers and clients in two districts of Ghana. Health Economics Review. 2012; 2:13.

8. Aryeetey GC, Jehu-Appiah C, Spaan E, Agyepong I, Baltussen R. Costs, equity, efficiency and feasibility of identifying the poor in Ghana’s National Health Insurance Scheme: empirical analysis of various strategies. Tropical Medicine and International Health. 2011;1365–3156.

9. National Health Insurance Authority (NHIA). National Health Insurance Scheme in Ghana: Reforms and Achievements. Accra: NHIA. 2013.

10. Witter S, Garshong B. Something old or something new? Social Health Insurance in Ghana. BMC International Health and Human Rights. 2009;9:20.

11. Sarpong N, Loag W, Fobil J, Meyer CG, Adu-Sakodie Y, May J, et al. National health insurance coverage and socio-economic status in a rural district of Ghana. Tropical Medicine and International Health. 2010;15(2):191–197.

12. Aryeetey GC, Jehu-Appiah C, Spaan E, D’Exelle B, Agyapong I, Baltussen R. Identification of poor households for premium exemptions in Ghana’s National Health Insurance Scheme: empirical analysis of three strategies. Tropical Medicine and International Health. 2010;1365–3156.

13. Macha J, Harris B, Grashong B, Ataguba JE, Akazili J, Kuwawenaruwa A, et al. Factors influencing the burden of health care financing and the distribution of health care benefits in Ghana, Tanzania and South Africa. Health Policy and Planning. 2012;27:i46–i54.

14. Fusheini A, Marnoch G, Gray AM. Stakeholders perspectives on the Success Drivers in Ghana’s National Health Insurance Scheme – Identifying Policy Translation Issues. Int. J Health Policy Manag. 2017;6(5):273–283.

15. Cooke E, Hague S, McKay A. The Ghana Poverty and Inequality Report. Using the 6th Ghana Living Standards. Available from: https://www.unicef.org/ghana/Ghana_Poverty_and_Inequality_Analysis_FINAL_Match_2016(1).pdf.

16. National Health Insurance Authority (NHIA). NHIA role and functions. Accra: NHIA. 2015. Available from: http://www.nhis.gov.gh/districts.aspx.

17. Agyepong IA, Abankwah DNY, Abroso A, Chun C, Dodoo JNO, Lee S, et al. National Health Insurance Scheme: policy and implementation challenges and dilemmas of a lower middle income country. BMC Health Services Research. 2016;16:504.

18. Awoonor-Williams JK, Tindana P, Dalinjong PA, Nartey H, Akazili J. Does the operations of the National Health Insurance Scheme (NHIS) in Ghana align with the goals of Primary Health Care? Perspectives of key stakeholders in northern Ghana. BMC International Health and Human Rights. 2016;16:21.

19. Ministry of Health (MOH). Holistic Assessment of the Health Sector Programme of Work 2017. Accra: MOH. 2018.

20. Gobah FKF, Liang Z. The National Health Insurance Scheme in Ghana: Prospects and Challenges; a Cross-Sectional Evidence. Global Journal of Health Science. 2011;1916–9736.

21. Schieber G, Cashin C, Saleh K, Lavado R. Health Financing in Ghana. Washington DC: World Bank. 2011. doi:10.1596/9780-8213-9566-0.

22. National Health Insurance Authority (NHIA). 2012 Annual Report. Accra: NHIA. 2013.

23. Alfers L. The Ghana National Health Insurance Scheme: Barriers to Access for Informal Workers. WEIGO Working Paper (Social Protection): No. 30. 2013.

24. National Health Insurance Authority (NHIA). 2011 Annual Report. Accra: NHIA. 2012.

25. National Health Insurance Authority (NHIA). 2013 Annual Report. Accra: NHIA. 2014.

26. Ministry of Health (MOH). Holistic Assessment of the Health Sector Programme of Work 2015. Accra: MOH. 2016.

27. World Health Organization (WHO). WHO Country Office for Ghana: Annual Report 2014. Accra: WHO. 2015.

28. Apoya P, Marriott A. Achieving a shared universal goal: free universal health Care in Ghana. Accra: OXFAM. 2011. Available from: http://policy-practice.oxfam.org.uk/publications/achieving-a-shared-goal-free-universalhealthcare-in-ghana-125306.

29. Seddoh A, Sataru F. Mundane? Demographic characteristics as predictors of enrolment onto the National Health Insurance Scheme in two districts of Ghana. BMC Health Services Research. 2018;18:330.

30. Huihui W, Otoo N, Dsane-Selby L. Ghana National Health Insurance Scheme: improving financial sustainability based on expenditure review. A World Bank study. Washington, D.C.: World Bank Group. 2017. https://doi.org/10.1596/978-1-4648-1117-3.

31. Alhassan RK, Duku SO, Janssens W, Nketiah-Amponsah E, Spieker N, van Ostenberg P, et al. Comparison of Perceived and Technical Healthcare Quality in Primary Health Facilities: Implications for a Sustainable National Health Insurance Scheme in Ghana. PLoSONE. 2015:10(10): e0140109. doi:10.1371/journal.pone.0140109.

32. Abuosi AA, Domfeh KA, Abor JY, Nketiah-Amponsah E. Health insurance and quality of care: Comparing perceptions of quality between insured and uninsured patients in Ghana’s hospitals. International Journal for Equity in Health. 2016;15:76.

33. Ghana Statistical Service (GSS). 2010 Population and Housing Census, District Analytical Report: Ashaiman Municipality. Accra: GSS. 2014.

34. Ghana Statistical Service (GSS). Poverty Profile in Ghana (2005–2013): Ghana Living Standards Survey Round 6. Accra: GSS. 2014.

35. African Health Markets for Equity (AHME). Monitoring and evaluation framework for the African Health Markets for Equity project in Ghana: scope of work. Accra: International Finance Corporation. 2015.

36. Ghana Statistical Service (GSS). 2010 Population and Housing Census, District Analytical Report: Adaklu-Anyigbe District. Accra: GSS. 2014.

37. Ghana Statistical Service (GSS). 2010 Population and Housing Census, District Analytical Report: Kasena Nankana District. Accra: GSS. 2014.

38. Nelson OC, Kalu O, Jimmy E, Uwanede CC, Abeshi SE, Offiong DA. Evaluating the impact of National Health Insurance Scheme on health care consumers in Calabar metropolis, Southern Nigeria. International Journal of Learning & Development. 2013:2164–4063.

39. Bai YK, Middlestadt SE, Peng CYJ, Flys AD. Psychosocial factors underlying the mother’s decision to continue exclusive breastfeeding for six months: an elicitation study. Journal of Human Nutrition and Dietetics. 2009;134–140.

